# Recent demographic histories and genetic diversity across pinnipeds are shaped by anthropogenic interactions and mediated by ecology and life-history

**DOI:** 10.1101/293894

**Authors:** M.A. Stoffel, E. Humble, K. Acevedo-Whitehouse, B.L. Chilvers, B. Dickerson, F. Galimberti, N. Gemmell, S.D Goldsworthy, H.J. Nichols, O Krüger, S. Negro, A. Osborne, A.J. Paijmans, T. Pastor, B.C. Robertson, S. Sanvito, J. Schultz, A.B.A Shafer, J.B.W. Wolf, J.I. Hoffman

## Abstract

A central paradigm in conservation biology is that population bottlenecks reduce genetic diversity and negatively impact population viability and adaptive potential. In an era of unprecedented biodiversity loss and climate change, understanding both the determinants and consequences of bottlenecks in wild populations is therefore an increasingly important challenge. However, as most studies have focused on single species, the multitude of potential drivers and the consequences of bottlenecks remain elusive. Here, we used a comparative approach by integrating genetic data from over 11,000 individuals of 30 pinniped species with demographic, ecological and life history data to elucidate the consequences of large-scale commercial exploitation by 18^th^ and 19^th^ century sealers. We show that around one third of these species exhibit strong genetic signatures of recent population declines, with estimated bottleneck effective population sizes reflecting just a few tens of surviving individuals in the most extreme cases. Bottleneck strength was strongly associated with both breeding habitat and mating system variation, and together with global abundance explained a large proportion of the variation in genetic diversity across species. Overall, there was no relationship between bottleneck intensity and IUCN status, although three of the four most heavily bottlenecked species are currently endangered. Our study reveals an unforeseen interplay between anthropogenic exploitation, ecology, life history and demographic declines, sheds new light on the determinants of genetic diversity, and is consistent with the notion that both genetic and demographic factors influence population viability.

## Introduction

Unravelling the demographic histories of species is a fundamental goal of population biology and has tremendous implications for understanding the genetic variability observed today ^1,2^. Of particular interest are sharp reductions in the effective population size (*N_e_*) known as population bottlenecks ^3,4^, which may negatively impact the viability and adaptive evolutionary potential of species through a variety of stochastic demographic processes and the loss of genetic diversity ^5–8^. Specifically, small bottlenecked populations have elevated levels of inbreeding and genetic drift, which decrease genetic variability and can lead to the fixation of mildly deleterious alleles and ultimately drive a vortex of extinction ^6,8–10^. Hence, investigating the bottleneck histories of wild populations and their determinants and consequences is more critical than ever before, as we live in an era where global anthropogenic alteration and destruction of natural habitats are driving species declines on an unprecedented scale ^11,12^.

Unfortunately, detailed information about past population declines across species is sparse because historical population size estimates are often either non-existent or highly uncertain ^13,14^. A versatile solution for inferring population bottlenecks from a single sample of individuals is to compare levels of observed and expected genetic diversity, the latter of which can be simulated under virtually any demographic scenario based on the coalescent ^15–17^. A variety of approaches based on this principle have been developed, one of the most widely used being the heterozygosity-excess test, which compares the heterozygosity of a panel of neutral genetic markers to the expectation in a stable population under mutation-drift equilibrium ^18^. Although theoretically well grounded, these methods are highly sensitive to the assumed mutation model, which is seldom known ^19^. A more sophisticated framework for inferring demographic histories is coalescent-based Approximate Bayesian Computation (ABC) ^20^. ABC has the compelling advantages of making it possible to (i) compare virtually any demographic scenario as long as it can be simulated, (ii) estimate key parameters of the model such as the bottleneck effective population size and (iii) incorporate uncertainty in the specification of models by using priors. Due to this flexibility, ABC has become a state of the art approach for inferring population bottlenecks as well as demographic histories in general ^20–26^.

Although the widespread availability of neutral molecular markers such as microsatellites has facilitated numerous genetic studies of bottlenecks in wild populations, the vast majority of studies focused exclusively on single species and were confined to testing for the presence or absence of bottlenecks. We therefore know very little about the intensity of demographic declines and how these are influenced by anthropogenic impacts as well as by factors intrinsic to a given species. For example, species occupying breeding habitats that are more accessible to humans would be expected to be at higher risk of declines, while species with highly skewed mating systems tend to have lower effective population sizes ^27^ and might also experience stronger demographic declines as only a fraction of individuals contribute towards the genetic makeup of subsequent generations. Consequently, in order to disentangle the forces shaping population bottlenecks, we need comparative studies incorporating genetic, ecological and life history data from multiple closely related species within a consistent analytical framework.

Another question that remains elusive due to a lack of comparative studies is to what extent recent bottlenecks have impacted the genetic diversity of wild populations. While a number of influential studies of heavily bottlenecked species have indeed found very low levels of genetic variability ^28–31^ others have reported unexpectedly high genetic variation after supposedly strong population declines ^24,32–34^. Hence, it is not yet clear how population size changes contribute towards one of the most fundamental questions in evolutionary genetics – how and why genetic diversity varies across species ^2,35–37^. To tackle this question, we need to compare closely related species because deeply divergent taxa vary so profoundly in their genetic diversity due to differences in their life-history strategies that any effects caused by variation in *N_e_* will be hard to detect and decipher ^36–37^.

Finally, the relative contributions of genetic diversity and demographic factors towards extinction risk remain unclear. While historically there has been a debate about the immediate importance of genetic factors towards species viability ^5,7^ there is now growing evidence that low genetic diversity increases extinction risk ^8,38^ and on a broader scale that threatened species show reduced diversity ^7^. Nevertheless, due to a lack of studies measuring bottlenecks consistently across species, it remains an open question as to how the loss of genetic diversity caused by demographic declines ultimately translates into a species extinction risk, which is assessed by its International Union for Conservation of Nature (IUCN) status.

An outstanding opportunity to address these questions is provided by the pinnipeds, a clade of marine carnivores inhabiting nearly all marine environments ranging from the poles to the tropics and showing remarkable variation in their ecological and life-history adaptations ^39^. Pinnipeds include some of the most extreme examples of commercial exploitation known to man, with several species including the northern elephant seal having been driven to the brink of extinction for their fur and blubber by 18^th^ to early 20^th^ century sealers ^13^. By contrast, other pinniped species inhabiting pristine environments such as Antarctica have probably had very little contact with humans ^13^. Hence, pinnipeds show large differences in their demographic histories within the highly constrained time window of commercial sealing and thereby represent a unique ‘natural experiment’ for exploring the causes and consequences of recent bottlenecks.

Here, we conducted a broad-scale comparative analysis of population bottlenecks using a combination of genetic, ecological and life-history data for 30 pinniped species. We inferred the strength of historical declines across species from the genetic data using two complimentary coalescent-based approaches, heterozygosity-excess and ABC. Heterozygosity-excess was used as a measure of the relative strength of recent population declines, while a consistent ABC framework was used to evaluate the probability of each species having experienced a severe bottleneck during the known timeframe of commercial exploitation, as well as to estimate relevant model parameters. Finally, we used Bayesian phylogenetic mixed models to investigate the potential causes and consequences of past bottlenecks while controlling for phylogenetic relatedness among species. We hypothesised that (i) extreme variation in the extent to which species were exploited by man should be reflected in their genetic bottleneck signatures; (ii) both breeding habitat and mating system should have an impact on the strength of bottleneck signatures across species; (iii) past bottlenecks should reduce contemporary genetic diversity; and (iv) heavily bottlenecked species with reduced genetic diversity will be more likely to be of conservation concern.

## Results

### Genetic data

We analysed a combination of published and newly generated microsatellite data from 30 pinniped species, with a median of 253 individuals and 14 loci per species (see Methods and Supplementary Table 1 for details). Measures of genetic diversity, standardised across datasets as the average per ten individuals, varied considerably across the pinniped phylogeny, with observed heterozygosity (*H_o_*) and allelic richness (*A_r_*) varying by over two and almost five-fold respectively across species (Supplementary Table 2). Both of these measures were highly correlated (*r* = 0.92) and tended to be higher in ice breeding seals, intermediate in fur seals and sea lions, and substantially lower in a handful of species including northern elephant seals and monk seals (Fig 1A).

**Fig. 1:**
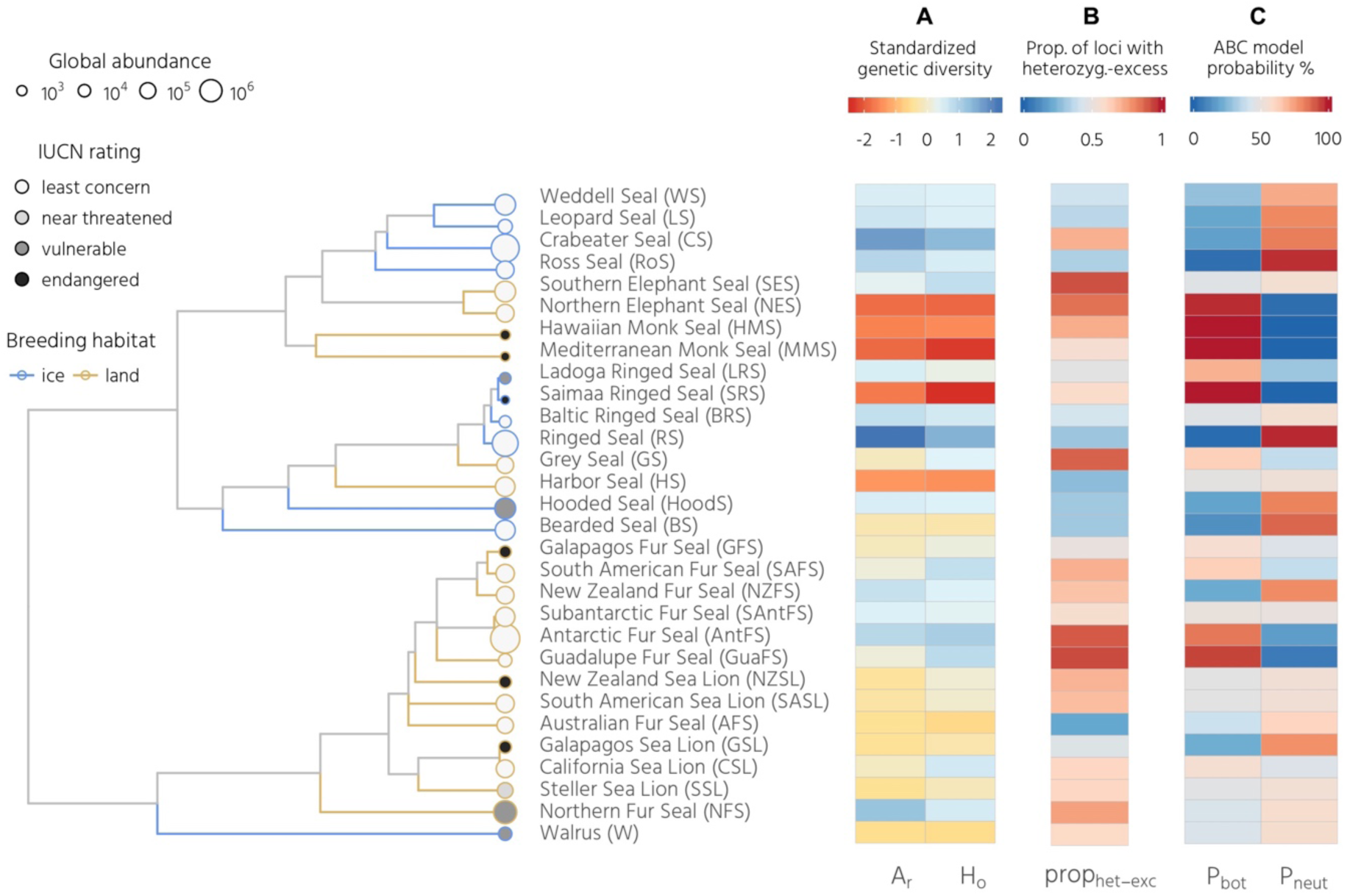
Patterns of genetic diversity and bottleneck signatures across the pinnipeds. The phylogeny shows 30 species with branches colour coded according to breeding habitat and tip points coloured and sized according to their IUCN status and global abundance respectively. Panel A shows two genetic diversity measures, allelic richness (*A*_*r*_) and observed heterozygosity (*H*_*o*_), standardized across species. Panel B shows the proportion of loci in heterozygosity-excess (*prop*_het-exc_) calculated for the TMP80 model (see Methods for details). Panel C summarises the ABC model selection results, with posterior probabilities corresponding to the bottleneck versus constant population size model. These data are also summarised in Supplementary Table 2 and 3.

### Bottleneck inference

We used two different coalescent-based approaches to infer the extent of recent population bottlenecks. First, the amount of heterozygosity-excess at selectively neutral loci such as microsatellites is an indicator of recent bottlenecks because during a population decline the number of alleles decreases faster than heterozygosity ^3^. Recent bottlenecks therefore generate a transient excess of heterozygosity relative to a population at equilibrium with an equivalent number of alleles ^18^. Here, we quantified the proportion of loci in heterozygosity-excess (*prop*_het-exc_) for each species, which was highly repeatable across a range of mutation models (see Methods and Supplementary Table 3). Consequently, we focused on a two-phase model with 80% single-step mutations (TPM80), which is broadly in line with mammalian mutation model estimates from the literature ^40^ as well as posterior estimates from our ABC analysis (Supplementary table 4). Fig. 1B shows a heatmap of *prop*_het-exc_ across species, which is bounded between zero (all loci show heterozygosity-deficiency, an indicator of recent expansion) and one (all loci show heterozygosity-excess, an indicator of recent decline) whereby 0.5 is the expectation for a stable population. Considerable heterogeneity was found across species, with northern and southern elephant seals, grey seals, Guadalupe fur seals and Antarctic fur seals showing the strongest bottlenecks signals. By contrast, the majority of ice-breeding seals showed heterozygosity-deficiency, consistent with historical population expansions.

Second, we used ABC to select between a bottleneck and a neutral model as well as to estimate posterior distributions of relevant parameters. To optimally capture recent population size changes across species, we allowed *N_e_* to vary over time in both models within realistic priors (see Methods) while the bottleneck model also included a severe decrease in *N_e_* to below 800 during the time of peak sealing. ABC was clearly able to distinguish between the two models, with the posterior probability of correct model classification being 75% for the bottleneck model and 71% for the neutral model, and for every species the preferred model showed a good fit to the data (all *p*-values > 0.05, Supplementary table 5). The posterior bottleneck model probability (*p*_bot_) varied substantially across species and was strongly but imperfectly correlated with *prop*_het-exc_ (posterior median and 95% credible intervals; *β* = 0.17 [0.06, 0.27], R^2^_marginal_ = 0.38 [0.07, 0.64], see Supplementary Fig. 3). For eleven species, the bottleneck model was supported with a higher probability than the neutral model (i.e. *p*_bot_ > 0.5, see Supplementary Table 3). Subsequent parameter estimation was therefore based on the bottleneck model for eleven species and on the neutral model for the other 19 species.

**Figure 3:**
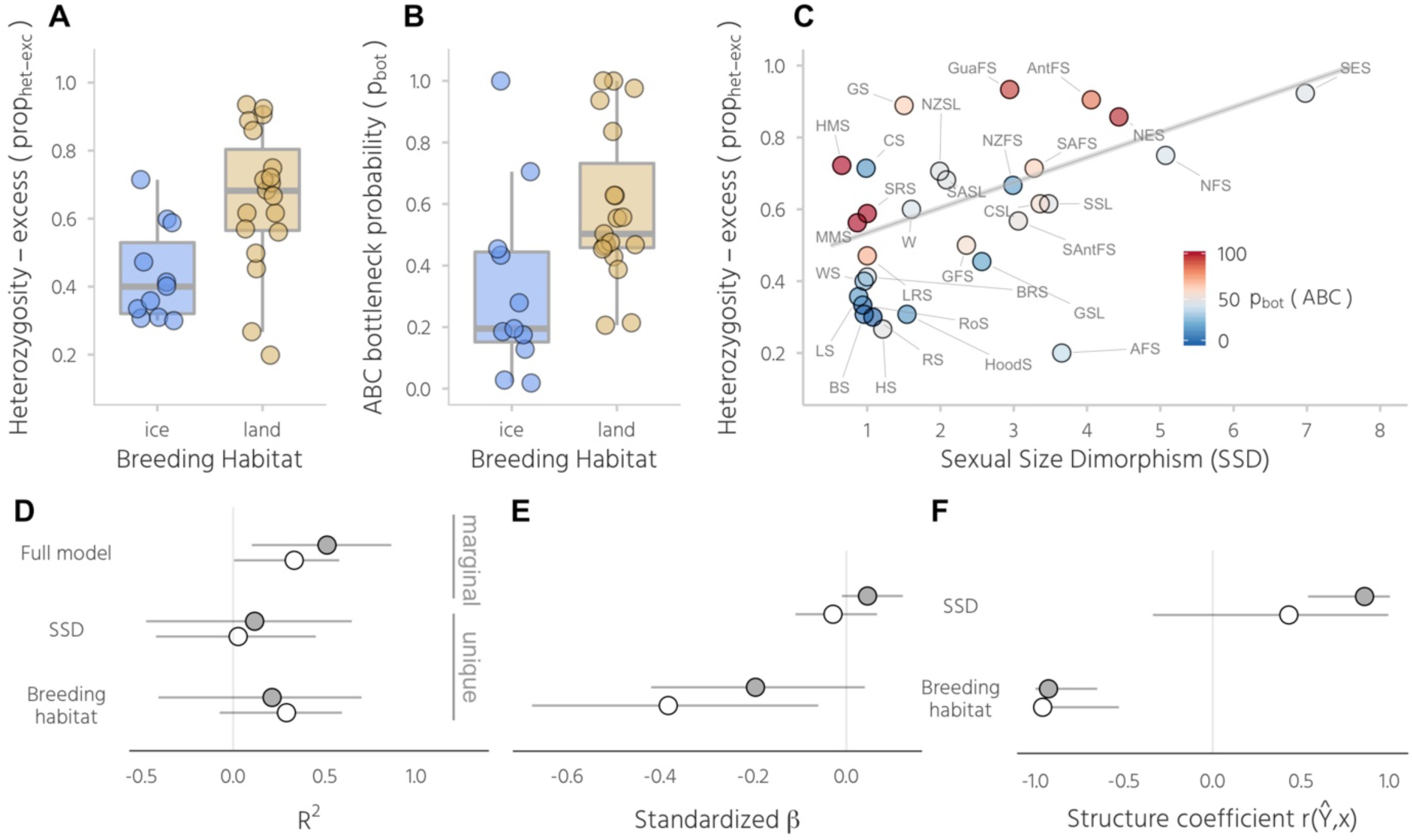
Ecological and life-history effects on bottleneck signatures. Shown are the results of phylogenetic mixed models of *prop*_het-exc_ and *p*_bot_ with breeding habitat and SSD fitted as fixed effects. Panels A and B show differences between ice- and land-breeding species in *prop*_het-exc_ and *p*_bot_ respectively. Raw data points are shown together with standard Tukey box plots. Panel C shows the relationship between sexual size dimorphism (SSD) and *prop*_het-exc_, with individual points colour coded according to the ABC bottleneck probability (*p*_bot_) and the line representing the predicted response from the *prop*_het-exc_ .model. Marginal and unique *R*^2^ values, standardized *β* coefficients and structure coefficients are shown for models of *prop*_het-exc_ (filled points) and *p*_bot_ (open points) in panels D-F, where they are presented as posterior medians with 95% credible intervals. Species abbreviations are given in Fig. 1 and Supplementary table 1.

Under the bottleneck model, prediction errors from the cross-validation were well below one for the bottleneck effective population size (*N_e_bot,* Supplementary Table 4A) and mutation rate (*µ*, Supplementary Table 4A) indicating that posterior estimates contain information about the underlying true parameter values. Similarly, under the neutral model, *µ* (Supplementary Table 4B) and the parameter describing the proportion of multi-step mutations (*GSM*_par_, Supplementary Table 4B) were informative. By contrast, although the pre-bottleneck effective population size (*N_e_hist*) also had a prediction error below one in both models, visual inspection of the cross-validation results revealed high variation in the estimates and a systematic underestimation of larger *N_e_hist* values, so this parameter was not considered further. Fig. 2 shows the eleven bottlenecked species ranked in descending order of estimated posterior modal *N*_e_*bot* (see also Supplementary Table 4A). The parameter estimates were indicative of strong bottlenecks (i.e. 200 < *N*_e_*bot* < 700) in seven species including both phocids and otariids, while even smaller *N*_e_*bot* values (i.e. *N*_e_*bot <* 100) were estimated for four phocids comprising the landlocked Saimaa ringed seal, both monk seal species and the northern elephant seal. Mutation rate estimates were remarkably consistent across species, with modes of the posterior distributions typically varying around 1 x 10^−4^ (Supplementary Fig. 1 and Supplementary Table 4), while *GSM*_par_ across species typically varied between around 0.2 and 0.3 (See Supplementary Fig. 2 and Supplementary Table 4B). Therefore, although studies of individual species are usually limited by uncertainty over the underlying mutation characteristics, our ABC analyses converged on similar estimates of mutation model and rate across species, allowing us to appropriately parameterise our bottleneck analyses.

**Figure 2:**
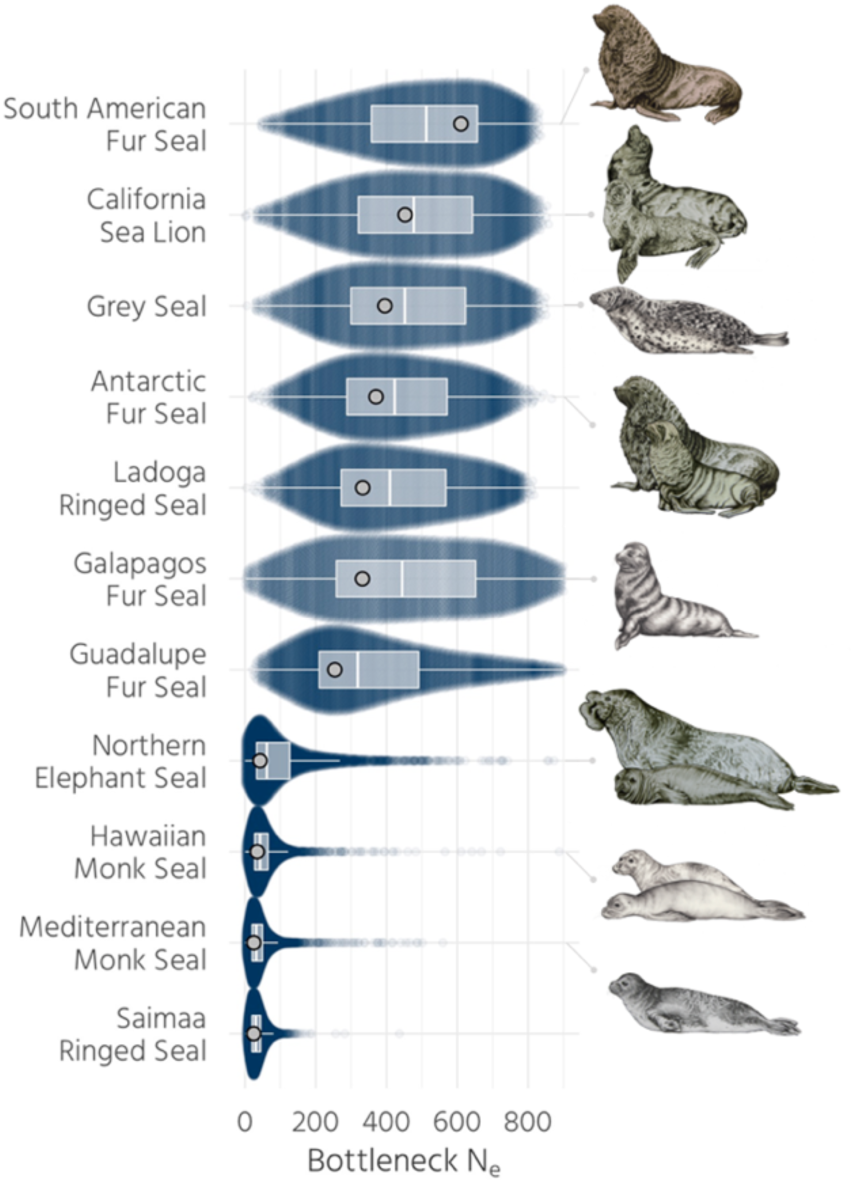
Estimated bottleneck effective population sizes. Posterior distributions of *N*_e_*bot* are shown for eleven species for which the bottleneck model was supported in the ABC analysis, ranked according to the modes of their density distributions which reflect the estimated most likely *N*_e_*bot.* Prior distributions are not shown as *N*_e_*bot* was drawn from a uniform distribution with U[1, 800]. For each species, parameter values for 5,000 accepted simulations are presented as a sinaplot, which arranges the data points to reflect the estimated posterior distribution. Superimposed are Tukey boxplots with light grey points representing maximum densities. (*Note to the editor and reviewers: The pinniped drawings are original artwork and the ringed seals and the Guadalupe fur seal are still pending and will be included when finished*)

To explore whether our results could be affected by population structure, we used STRUCTURE ^41^ to infer the most likely number of genetic clusters (*K*) across all datasets. For all of the species for which the best supported value of *K* was more than one (*n* = 12), we recalculated genetic summary statistics and repeated the bottleneck analyses based on individuals comprising the largest cluster. Using the largest genetic clusters did not appreciably affect our results, with repeatabilities for the genetic summary statistics and bottleneck signatures all being greater than 0.9 (see Supplementary table 6 for repeatabilities1 and Supplementary Fig. 3, which is virtually identical to Fig. 1).

### Factors affecting bottleneck history

Conceivably both ecological and life-history variables could have impacted the extent to which commercial exploitation affected different pinniped species. We hypothesised that breeding habitat would be an important ecological variable, as ice-breeding species are less accessible and more widely dispersed than their land breeding counterparts. Furthermore, sexual size dimorphism (SSD) should be an important life history variable, as species with a high SSD aggregate in denser breeding colonies making them more valuable to hunters, and polygyny reduces effective population size. We found clear differences between ice- and land-breeding seals in both *prop*_het-exc_ and *p_bot_*, with land-breeders on average showing stronger bottleneck signatures (Figs 3A,B). In addition, *prop*_het-exc_ was positively associated with SSD but no clear relationship was found with *p*_bot_ (Fig. 3C). To investigate this further, we constructed two Bayesian phylogenetic mixed models with *prop*_het-exc_ and *p*_bot_ as response variables respectively and SSD and breeding habitat fitted as predictors (see Methods for details). Both models explained an appreciable amount of variation (*prop*_het-exc_ R^2^_marginal_ = 0.51, CI [0.10, 0.86];*p*_bot_ R^2^_marginal_ = 0.33, CI [0.005, 0.58], Fig. 3D). As breeding habitat and SSD are correlated (R^2^_marginal_ = 0.40, CI [0, 0.83]), we reported both standardised model estimates (*β*) and structure coefficients 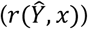, which represent the correlation between each predictor and the fitted response independent of the other predictors. Breeding habitat showed larger effect sizes than SSD in both models, although the corresponding credible intervals were also broader, indicating greater uncertainty (Fig 3E, Supplementary Table 8). By contrast, structure coefficients showed that breeding habitat and SSD were both strongly correlated to the fitted response in the *prop*_het-exc_ model, while SSD indeed had a much weaker effect in the *p*_bot_ model (Fig 3F, Supplementary Table 8). Thus, breeding habitat and SSD explain variation in *prop*_het-exc_ whereas only breeding habitat explains variation in *p*_bot_.

### Determinants of genetic diversity

To investigate the determinants of contemporary genetic diversity across pinnipeds, we constructed a phylogenetic mixed model of allelic richness (*A_r_*) with log transformed global abundance, breeding habitat and SSD fitted as predictor variables together with the two bottleneck measures *prop*_het-exc_ and \ (Fig. 4). A substantial 77% of the total variation in *A_r_* was explained (Fig. 4C, R^2^_marginal_ =0.77, CI [0.54, 0.93]). Specifically, *A_r_* increased by nearly five-fold from the least to the most abundant species (β = 1.26, CI [0.05, 2.43], Fig 4A), decreased nearly five-fold from the species with the lowest *p*_bot_ to the species with the highest *p*_bot_ (*β* = −2.00, CI [−3.35, −0.74] Fig 4A) and was on average 27% higher in ice than in land-breeding seals (*β* = 1.70, CI [0.31, 3.29], Fig. 4A). Due to multicollinearity among the five predictor variables (Supplementary Table 7), standardized *β* estimates (Fig 4D) can be hard to interpret because of potential suppression effects ^42^. This is reflected in the low unique *R*^2^ values of the predictors relative to the marginal *R*^2^ of the full model (Fig. 4C). However, the structure coefficients (Fig. 4E) also revealed strong associations between the fitted model response and breeding habitat 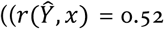, CI [0.24, 0.79]), abundance 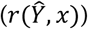 = 0.74, CI [0.55, 0.91]) and *p*_bot_ 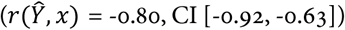 showing that all three variables are associated with the response.

**Figure 4:**
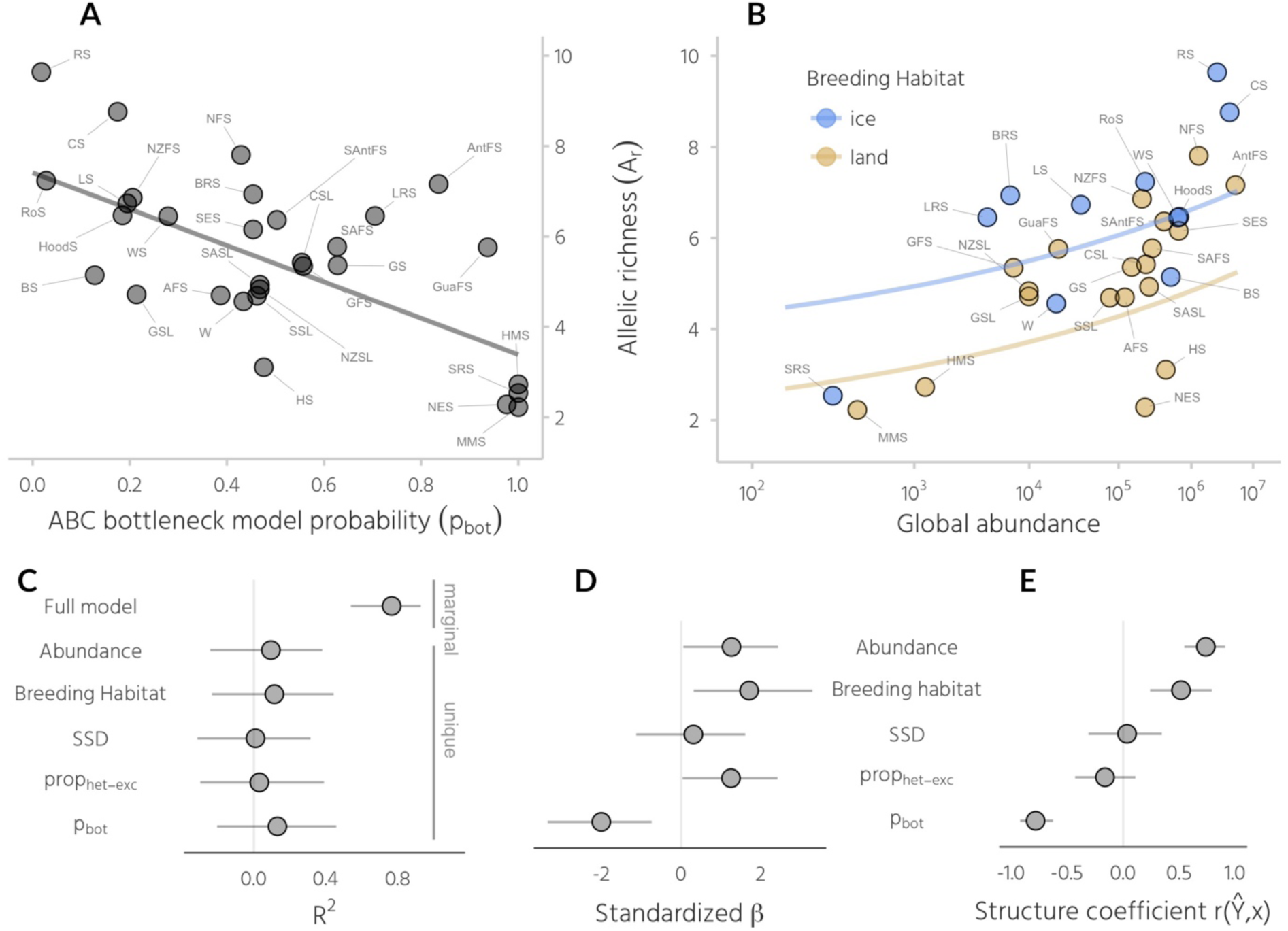
Determinants of current genetic diversity across pinnipeds. Panel A shows the relationship between global abundance and allelic richness (*A_r_*) with the grey line representing the model prediction. Panel B shows a scatterplot of *A_r_* versus *p*_bot_ with the lines representing model predictions for ice- and land-breeding seals respectively. Marginal and unique *R*^2^ values, standardised *β* estimates and structure coefficients for the model are shown respectively in panels C-E, where they are presented as posterior medians with 95% credible intervals. Species abbreviations are given in Fig. 1 and Supplementary table S1.

### Conservation status, bottleneck signatures and genetic diversity

To investigate whether population bottlenecks and low genetic diversity are detrimental to species viability, we asked whether contemporary conservation status is related to both the strength of past bottlenecks and to *A*_r_. Based on data from the IUCN red list ^43^, we classified species into two groups; the first comprised species listed as ‘*least concern*’ while the second combined species listed as ‘*near threatened’*, *’vulnerable*’ or ‘*endangered*’ into a ‘*concern*’ category. Using a phylogenetic mixed model, we did not find any clear differences in either heterozygosity-excess or *p*_bot_ with respect to conservation status (Fig. 6A, B). By contrast, average *A*_r_ was around 1.3 alleles lower in the *concern* category, although there was some uncertainty with the 95% credible interval of *β* ranging from −0.07 to 2.52.

**Figure 6:**
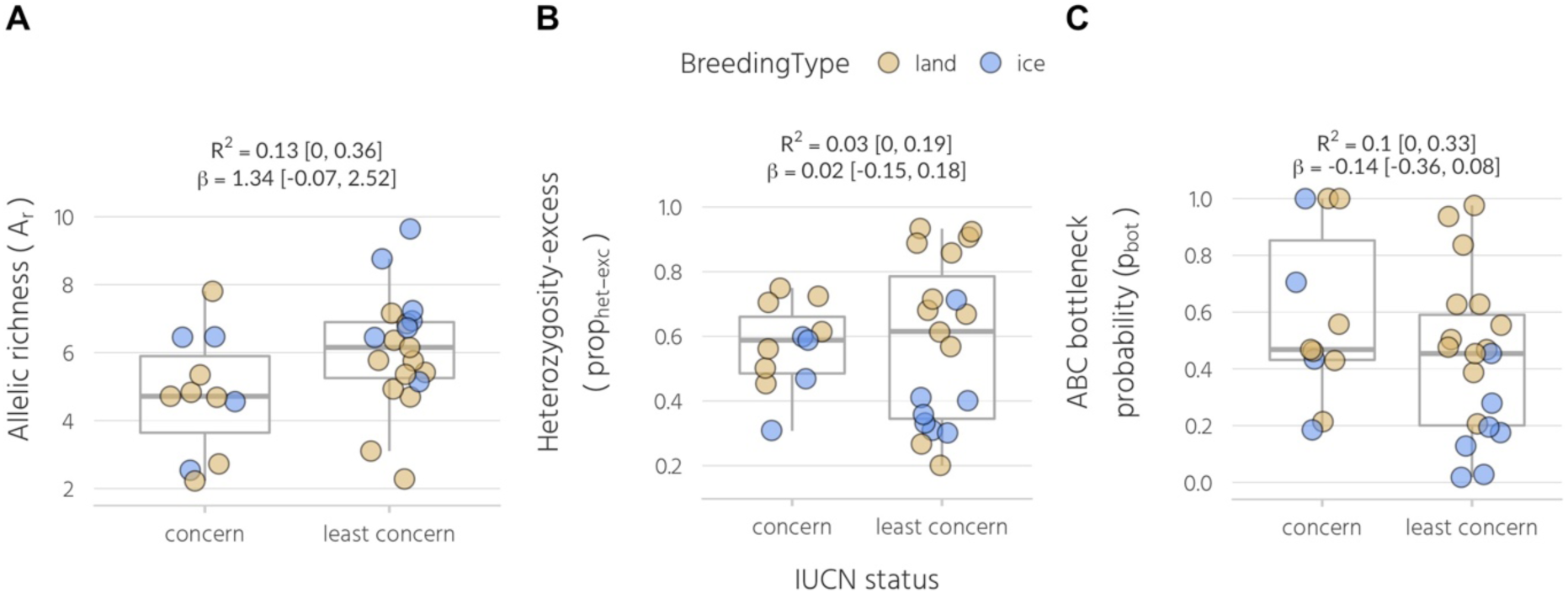
Conservation implications of bottlenecks and genetic diversity. All pinniped species were classified into either a *concern* or *least concern* category depending on their current IUCN status as described in the main text. Shown are the raw data for each category together with standard Tukey box plots for (A) *A*_r,_ (B) *prop*_het-exc_ and (C) *p*_bot_. Marginal *R*^2^ and standardised *β* estimates are shown for Bayesian phylogenetic mixed models with standardized predictors (see Materials and methods for details).

## Discussion

To explore the interplay between historical demography, ecological and life-history variation, genetic diversity and conservation status, we used a comparative approach based on genetic data from over 80% of all extant pinniped species. To model bottleneck strength, we used two approaches that capture different but complementary facets of genetic diversity resulting from population bottlenecks. Using ABC, we contrasted a bottleneck model incorporating a severe decrease in *N*_e_ during the time of peak sealing in the 18^th^ and 19^th^ Centuries with a neutral model. The resulting bottleneck measure, *p*_bot_ is the probability (relative to the neutral model) that a species’ observed genetic diversity is similar to the diversity of a population that experienced a severe reduction in *N*_e_ below 800, and therefore provides an *absolute* bottleneck measure. By contrast, heterozygosity excess (*prop*_het-exc_) theoretically captures sudden recent reductions in *N*_e_ even in fairly large populations ^18^, and therefore provides a *relative* bottleneck measure. Concretely, given the average sample size of individuals and loci used in this study, we would expect to detect an excess of heterozygosity at the majority of loci (i.e. *prop*_het-exc_ > 0.5) when a 100− to 1000−fold reduction in *N*_e_ occurred within the last 4*N*_e_ generations, regardless of the magnitude of *N*_e_ (see simulations in ^18^).

ABC analysis supported the bottleneck model for more than a third of the species. The strongest bottlenecks (*N*_e_*bot* <50) were inferred for the northern elephant seal, a textbook example of a species that bounced back from the brink of extinction ^44^, as well as for the two monk seals and the Saimaa ringed seal, species with very small geographic ranges and a long history of anthropogenic interaction ^13^. Slightly weaker bottlenecks were estimated for seven further species including Antarctic and Guadalupe fur seals, both of which share a known history of commercial exploitation for their fur ^13^. At the other end of the continuum, several Antarctic species that have not been commercially hunted such as crabeater and Weddell seals showed unequivocal support for the neutral model in line with expectations. Surprisingly, several otariid species known to have been hunted in the hundreds of thousands (e.g. South American sea lions) to millions (e.g. northern fur seals) did not show clear support for a bottleneck as strong as simulated in our analyses. This suggests that sufficiently large numbers of individuals must have survived despite extensive sealing, possibly on inaccessible shores or remote islands ^45^.

We hypothesised that not all pinniped species were equally affected by commercial exploitation partly due to intrinsic differences relating to a species’ ecology and life-history. In line with this, we found a strong influence of breeding habitat on bottleneck signatures, with both *prop*_het-exc_ and *p*_bot_ being higher in species that breed on land relative to those breeding on ice. A likely reason for this is that terrestrially breeding pinniped species were more profitable due to their generally higher population densities and accessibility, and therefore probably experienced more intense hunting. We also found that heterozygosity-excess was strongly linked to sexual size dimorphism (SSD), with highly polygynous species like elephant seals and some fur seals showing the strongest footprints of recent decline. While this could reflect the increased ease of exploitation and thus higher commercial value of species that predictably aggregate in very large numbers to breed, species with higher SSD also have highly skewed mating systems making them potentially more vulnerable to severe decreases in *N*_e_ when key males are taken out of the system. By contrast, we did not find an effect of SSD on the ABC bottleneck probability *p*_bot_, suggesting that although sexually dimorphic species experienced the greatest declines, these were not necessarily as severe as simulated in the ABC analysis (*N*_e_ < 800). This is probably because many species reached ‘economic extinction’ well above this threshold, when populations became too small to sustain the sealing industry.

Although vast numbers of species are declining globally at unprecedented rates ^12^ we still lack a clear understanding of how recent declines in *N*_e_ affect contemporary genetic diversity in wild populations ^2,36^. Here, we explained a large proportion of the five-fold variation in allelic richness (*A*_r_) observed from the most to the least diverse pinniped species. First, *A*_r_ was strongly associated with *p*_bot_ but not with *prop*_het-exc_, in agreement with the theoretical expectation that populations have to decline to a very small *N*_e_ ^3^, as was simulated in our ABC analysis, to lose a substantial proportion of their diversity. Second, we showed that global abundance across species was tightly linked to *A*_r_, despite the likely impact of bottlenecks and the limited time-window for the recovery of genetic diversity. As differences in genetic diversity across species are largely determined by long-term *N*_e_ ^2^, this implies that contemporary population sizes across pinnipeds must to some extent resemble patterns of historical abundance, and hence that many bottlenecked species have to a large extent rebounded to occupy their original niches. Third, *A*_r_ was higher in ice-breeding relative to land-breeding seals. However, a low unique *R*^2^ of breeding habitat in our model suggests that this probably reflects the more intense bottleneck histories of land-breeding seals rather than a true ecological effect.

Finally, we compared genetic diversity and bottleneck strength between species that are currently classified by the IUCN as being of conservation concern versus those that are not. We found that *A*_r_ was on average around 22% lower in species within the *concern* category, consistent with previous evidence from a broad range of species ^7^. While three out of the four pinniped species with the strongest estimated bottlenecks are currently listed as endangered, species from both categories did not overall differ in their bottleneck signatures. Our comparative study of population bottlenecks is therefore encouraging: population bottlenecks do not necessarily result in reduced genetic diversity, population viability and adaptive potential. As shown here, global bans on commercial sealing at the beginning of the 20^th^ Century allowed many surviving pinniped populations to recover in abundance. Those that have not sufficiently rebounded illustrate the two fundamental conservation challenges, especially as biodiversity loss and climate change continue at unprecedented rates: halting population declines and promoting population recovery.

## Methods

### Genetic data

Microsatellite data were obtained from a total of 30 pinniped species including three subspecies of ringed seal (summarised in Supplementary Table 1). We generated new data for five of these species as described in the Supplementary methods, while the remaining datasets were previously published. Sample sizes of individuals ranged between 16 for the Ladoga ringed seal to 2386 for the Hawaiian monk seal, with a median of 253 individuals. The number of loci genotyped varied between five and 35 with a median of 14.

### Phylogenetic, demographic, life history and conservation status data

Phylogenetic data were downloaded from the 10k trees website ^46^ and plotted using ggtree ^47^. The three ringed seal subspecies were added according to their separation after the last ice age ^48^. Demographic and life-history data for each species were obtained from ^49^. While most data stayed untransformed, we calculated sexual size dimorphism (SSD) as the ratio of male to female body mass, and log-transformed abundance across species to account for the several orders of magnitude differences in population sizes. Data on conservation status were retrieved from the IUCN website (http://www.iucnredlist.org/, 2017) ^43^.

### Data cleaning and preliminary population genetic analyses

In order to maximise data quality, we checked all datasets by eye and generated summary statistics and tables of allele counts to identify potentially erroneous genotypes including typographical or formatting errors. In ambiguous cases, we contacted the authors to verify the correct genotypes. As several of the datasets included samples from more than one geographical location, we used a Bayesian approach implemented in STRUCTURE version 2.3.4 ^41^ to infer the most likely number of genetic clusters (*K*) across all datasets. For computational and practical reasons, we used the ParallelStructure package in R ^50^ to run these analyses on a computer cluster. For all of the species for which the best supported value of *K* was more than one, we recalculated genetic summary statistics and repeated the bottleneck analyses based on individuals comprising the largest cluster and calculated repeatabilities including confidence intervals for all variables using the rptGaussian function in the rptR package ^51^. We also tested all loci from each dataset for deviations from Hardy-Weinberg equilibrium (HWE) using *χ*^2^ and exact tests implemented in pegas ^52^ and applied Bonferroni correction to the resulting *p*-values. Overall, 6% of loci were found to deviate from HWE in both tests and as these are unlikely to affect our comparative analyses, we focused subsequently on the full datasets.

### Genetic diversity statistics

In order to examine patterns of genetic diversity across species, we calculated observed heterozygosity (*H_o_*) and allelic richness (*A*_r_) with strataG {Archer:2017vj} as well as the proportion of low frequency alleles (*LFA*), defined as alleles with a frequency of <5%, using self-written code. For maximal comparability across species with different sample sizes, we randomly sampled ten individuals from each dataset 1000 times with replacement and calculated the corresponding mean and 95% confidence interval (CI) for each summary statistic.

### Heterozygosity-excess

We quantified heterozygosity-excess using the approach in ^18^ implemented in the program BOTTLENECK version 1.2.02 ^53^. BOTTLENECK compares the heterozygosity of a locus in an empirical sample to the heterozygosity expected in a population under mutation-drift equilibrium with the same number of alleles as simulated under the coalescent ^15,16^. Microsatellites evolve mainly by gaining or losing a single repeat unit ^54^ (the Stepwise Mutation Model, SMM), but occasional larger ‘jump’ mutations of several repeat units are also common ^55^. Consequently, BOTTLENECK allows the user to specify a range of mutation models, from the strict SMM through two-phase models (TPMs) with varying proportions of multi-step mutations to the infinite alleles model (IAM) where every new mutation is novel. We therefore evaluated the SMM plus three TPM models with 70%, 80% and 90% single-step mutations respectively and the default variance of the geometric distribution (0.30). For each of the mutational models, the heterozygosity of each locus expected under mutation-drift equilibrium given the observed number of alleles (*H_eq_*) was determined using 10000 coalescent simulations. The proportion of loci for which *H_e_* was greater than *H_eq_* (*prop*_het-exc_) was then quantified for all of the mutation models. To quantify consistency of the measure across mutation models, we calculated the repeatability of *prop*_het-exc_ using the rptR package ^51^ in R with 1000 bootstraps while adjusting for the mutation model as a fixed effect. Although the relative pattern across species was very consistent across mutation models (repeatability = 0.81, CI = [0.71, 0.89],), absolute values of *prop*_het-exc_ within species decreased with lower proportions of multistep mutations (means for the TPM70, 80, 90 and SMM were 0.63, 0.58, 0.49 and 0.27, respectively). Based on our posterior estimates (Supplementary Fig. 2) and in line with previous studies, we therefore based our subsequent analyses on *prop*_het-exc_ from the intermediate TPM80 model.

### Demographic models

As a second route to inferring historical population declines, we contrasted two alternative demographic scenarios (Fig. 7) using a coalescent-based approximate Bayesian computation (ABC) framework ^15,20,56,57^. To address the hypothesis that commercial exploitation from the 18^th^ to the beginning of the 20^th^ century led to population bottlenecks, we first defined a bottleneck model, which incorporates a severe reduction in population size within strictly bound time priors reflecting the respective time period. For comparison, we defined a neutral model, which did not contain a bottleneck, although it allowed the population size to vary over time within a defined set of priors. Genetic data under both models were simulated from broad enough prior distributions to fit all 30 species while keeping the priors as tightly bound as possible around plausible values. The bottleneck model was defined with seven different parameters (Fig. 7a). The current effective population size *N_e_* and the historical (i.e. pre-bottleneck) effective population size *N_e_hist* were drawn from a log-normal distribution with *N*_e_ ~ lognorm[logmean = 10.5, logsd = 1] and *N*_e_*hist* ~ lognorm[logmean = 10.5, logsd = 1]. This concentrated sampling within plausible ranges that fitted most species (i.e. with effective population sizes ranging from thousands to tens of thousands of individuals) while also occasionally drawing samples in the hundreds of thousands to fit the few species with very large populations. The bottleneck effective population size *N*_e_*bot* was drawn from a uniform distribution between 1 and 800 (*N*_e_*bot* ~ U[1, 800]) while the bottleneck start and end times *t_bot_start* and *t_bot_end* were drawn from uniform distributions ranging between ten and 70 (*t_bot_start* ~U[10, 70]) and one and 30 (*t_bot_end* ~ U[1, 30]) generations ago respectively. Hence, the bottleneck time priors encompassed the last four centuries for all species, as their estimated generation times vary between approximately 7 and 19 years (Supplementary Table 1). The microsatellite mutation rate *µ* was refined after initial exploration and drawn from a uniform prior with *µ* ~ U[10^−5^, 10^−4^] which lies within the range of current empirical estimates ^40,58^. The mutation model was defined as a generalized stepwise mutation model with the geometric parameter *GSM*_par_ reflecting the proportion of multistep mutations, uniformly distributed from *GSM_par_* ~U[0, 0.3]. The neutral model was defined with five parameters (Fig. 7b). *N*_e_, *N*_e_*hist, µ* and *GSM*_par_ were specified with the same priors as previously defined for the bottleneck model and the time parameter corresponding to the historical population size *t_hist_* was drawn from a uniform distribution ranging between 10 and 70 generations ago (*t_hist_* ~U[10, 70]).

**Figure 7:**
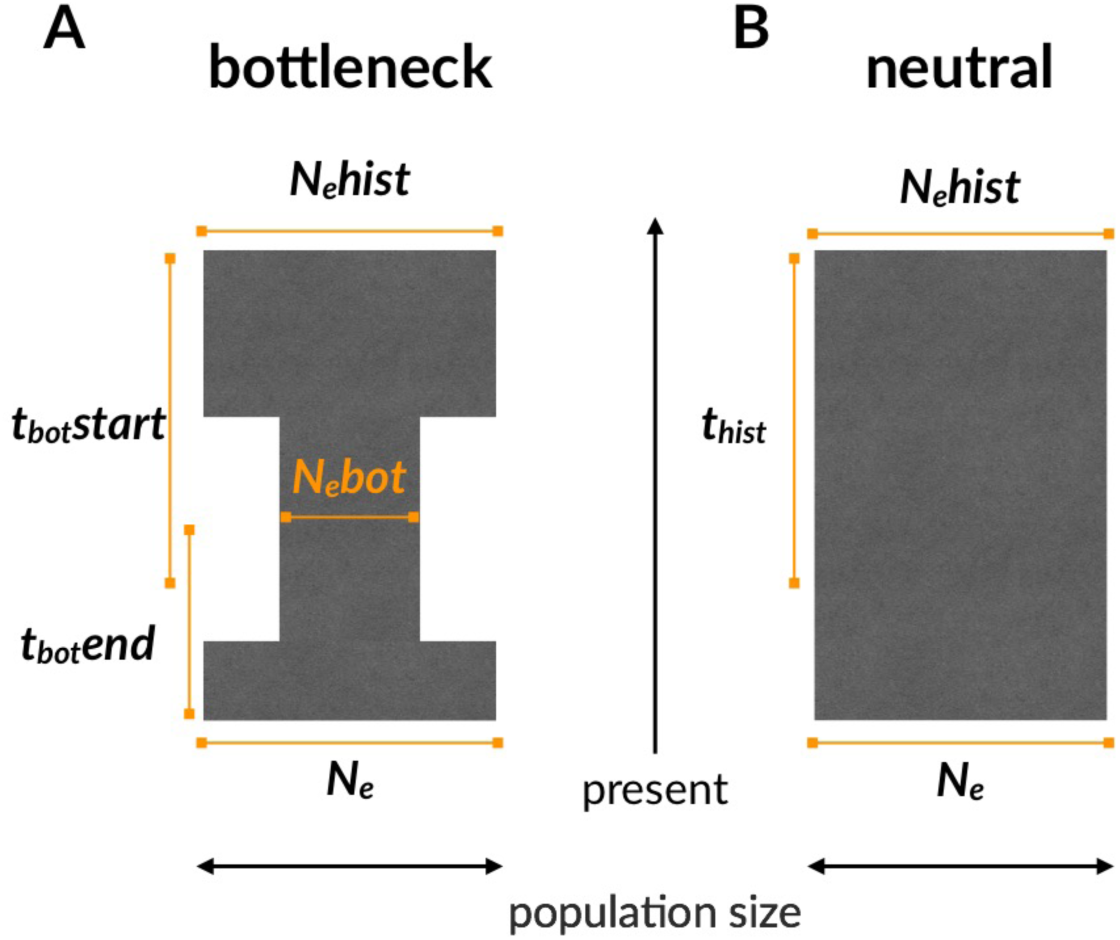
Schematic representation of two contrasting demographic scenarios and the parameter priors defining the models. The exact priors and mutation model are given in the Methods.

### ABC analysis

We simulated a total of 2 x 10^7^ datasets of 40 individuals and 10 microsatellite loci each under the two demographic scenarios using the fastsimcoal function in strataG ^59^ as an R interface to fastsimcoal2 ^60^, a continuous-time coalescent simulator. For both the simulated and empirical data, we used five different summary statistics for the ABC inference, all calculated as the mean across loci. Allelic richness (number of alleles), allelic range, expected heterozygosity (i.e. Nei’s gene diversity ^61^), the M-ratio ^62^ and the proportion of low frequency alleles (i.e. with frequencies < 5%). The summary statistics for the empirical datasets were computed by repeatedly re-sampling 40 individuals with replacement from the full datasets and calculating the mean across 1000 subsamples (for the Ladoga ringed seal and the Baltic ringed seal which had sample sizes smaller than 40, the full datasets were taken). As a small number of loci in the empirical data exhibited slight deviations from constant repeat patterns (i.e. not all of the alleles within a locus conformed to a perfect two, three or four bp periodicity), we calculated the M-ratio as an approximation using the most common repeat pattern of a locus to calculate the range of the allele size *r* and subsequently the M-ratio with M = *k*/(*r* + 1) where *k* is the number of alleles. All statistics were calculated using a combination of functions from the strataG package and self-written code. For the ABC analysis, we used a tolerance threshold of 5 x 10^−4^, thereby retaining 5000 simulations with summary statistics closest to those of each empirical dataset. For estimating the posterior probability for each scenario and each species, we used the multinomial regression method ^20,63^ as implemented in the function postpr in the abc package ^26^ where the model indicator is the response variable of a polychotomous regression and the accepted summary statistics are the predictors. To construct posterior distributions from the accepted summary statistics for the model parameters, we a local linear regression approach ^20^ as implemented in the abc function of the abc package.

### Evaluation of model specification and model fit via cross-validation

We evaluated whether ABC can distinguish between our two models by performing a leave-one-out cross validation implemented by the cv4postpr function of the abc package. Here, the summary statistics of one of the existing 2 x 10^7^ simulations were considered as pseudo-observed data and classified into either the bottleneck or the neutral model using all of the remaining simulations. If the summary statistics can discriminate between the models, a large posterior probability should be assigned to the model that generated the pseudo-observed dataset. This was repeated 100 times and the resulting posterior probabilities for a given model were averaged to derive the rate of misclassification. We furthermore checked for each species that the preferred model for each species provided a good fit to the empirical data by conducting a formal hypothesis test using the gfit function in abc. Specifically, we used the median distance between the accepted and observed summary statistics as a test statistic, whereby the null distribution was generated using summary statistics from the pseudo-observed datasets. Hence, a non-significant *p*-value indicates that the distance between the observed summary statistics and the accepted summary statistics is not larger than the expectation based on pseudo-observed data sets, i.e. the assigned model provides a good fit to the observed data.

### Evaluation of the accuracy of parameter estimates via cross-validation

In order to determine which parameters (i.e. population sizes, times and mutation rates and models) could be reliably estimated, we used leave-one-out cross validation implemented in the cv4abc function from the abc package to determine the accuracy of our ABC parameter estimates. For a randomly selected pseudo-observed dataset, parameters were estimated via ABC based on the remaining simulations using the rejection algorithm and a prediction error was calculated. This is possible because we know the “true” parameter values from which a given pseudo-observed dataset was simulated. This procedure was repeated 1000 times and a mean prediction error ranging between 0 and 1 was calculated, where 0 reflects perfect estimation and 1 means that the posterior estimate does not contain any information about the true parameter value ^26^.

### Bayesian phylogenetic mixed models

Finally, we used Bayesian phylogenetic mixed models in MCMCglmm ^64^ to evaluate the ecological and life-history variables affecting bottleneck strength and genetic diversity, and to test whether bottleneck history and genetic diversity are predictive of contemporary conservation status. Details of all the models are given in the supplementary material. All of the response variables were modelled with Gaussian distributions, while the predictors were fitted as fixed effects and the phylogenetic covariance matrix as a random effect. Predictors in models containing binary fixed effects were standardised by two standard deviations to allow a direct comparison between the effect sizes ^65^. In models without binary fixed effects, the predictor variables were standardised by one standard deviation. For all models, we report the marginal *R*^2^ as in ^66^. Some of the predictors in our models were inter-correlated and multicollinearity might lead to suppression effects and make the interpretation of regression coefficients difficult ^42^. We therefore reported standardized *β* estimates, structure coefficients, 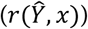 and unique *R*^2^ values for all variables in all models. The structure coefficients represent the correlation between a predictor and the fitted response of a model independent of the other predictors, and therefore reflect the direct contribution of a variable to that model. On the other hand, the unique *R*^2^ is the difference between the marginal *R*^2^ of a model including and a model excluding a predictor, which will be small when another predictor explains much of the same variation in the response ^42^. All model estimates were presented as the posterior median and 95% credible intervals (CIs). We used uninformative priors with a belief (shape) parameter v = 1 for the variance-covariance matrices of the random effects and inverse-Wishart priors with v = 0.002 for residual variances. For each model, three independent MCMC chains were run for 1,100,000 iterations, with a burn-in of 100,000 iterations and a thinning interval of 1000 iterations. Convergence was checked visually and by applying the Gelman-Rubin criterion to three independent chains. All of the upper 95% confidence limits of the potential scale inflation factors were below 1.05.

### Data and code availability

All data wrangling steps and statistical analyses except for the heterozygosity-excess tests ^53^ were implemented in R ^67^. The documented analysis pipeline along with the raw data can be accessed via GitHub (https://github.com/mastoffel/pinniped_bottlenecks) and is fully reproducible.

## Acknowledgements

This research was supported by standard grants (HO 5122/3-1 and HO 5122/5-1) from the German Research Foundation (DFG) and as part of the SFB TRR 212 (NC^3^) together with a dual PhD studentship from Liverpool John Moores University. We are grateful to John Arnould, David Coltman, Corey Davis, Larissa Rosa de Oliveira, Tom Gelatt and NOAA, Melissa Gladstone, Roger Kirkwood, Melanie Lancaster, Tim Malloy, Rolf Ream, Garry Stenson and Rob Stewart for providing access to published microsatellite datasets. We also thank Matthias Galipaud for advice on statistical analysis, Kevin Arbuckle for advice on Bayesian phylogenetic mixed models and Luke Eberhart-Phillips for helpful comments on the figures. Finally, we are very grateful to Rebecca Carter (www.rebeccacarterart.co.uk) for the time and effort she dedicated to producing the pinniped illustrations.

## Author contributions

Conceived the study: JIH and MAS. Generated data: KA, BLC, BD, FG, NG, SDG, HJN, OK, SN, AO, AJP, TP, BCR, SS, JS, ABAS, JBWW, JIH. Analysed data: MAS, EH and JIH. Wrote the paper: MAS and JIH. All of the authors commented upon and approved the final manuscript.

